# Agent-Based Modeling Demonstrates How Local Chemotactic Behavior Can Shape Biofilm Architecture

**DOI:** 10.1101/421610

**Authors:** Emily G. Sweeney, Andrew Nishida, Alexandra Weston, Maria S. Bañuelos, Kristin Potter, John Conery, Karen Guillemin

## Abstract

Mature bacterial biofilms have elaborate three-dimensional architectures that endow these structures with their durability and resistance to environmental perturbations. We used agent-based modeling to explore whether local cellular interactions were sufficient to give rise to global structural features of biofilms. Specifically, we asked whether chemorepulsion from a self-produced quorum-sensing molecule, autoinducer-2 (AI-2), was sufficient to recapitulate biofilm growth and cellular organization observed for biofilms of the human pathogen *Helicobacter pylori*. To carry out this modeling, we modified an existing platform, Individual-based Dynamics of Microbial Communities Simulator (iDynoMiCS), to incorporate three-dimensional chemotaxis, planktonic cells that could join or leave the biofilm structure, and cellular production of AI-2. We simulated biofilm growth of previously characterized *H. pylori* strains with varying AI-2 production and sensing capacities. Using biologically plausible parameters, we were able to recapitulate both the variation in biofilm mass and cellular distributions observed with these strains. Specifically, the strains that were competent to chemotax away from AI-2 produced smaller and more heterogeneously spaced biofilms, whereas the AI-2 chemotaxis defective strains produced larger and more homogeneously spaced biofilms. The model also provided new insights into the cellular demographics contributing to the biofilm patterning of each strain. Our analysis supports the idea that cellular interactions at small spatial and temporal scales are sufficient to give rise to larger scale emergent properties of biofilms.

**Importance:** Most bacteria exist in aggregated, three-dimensional structures called biofilms. Biofilms are resistant to antimicrobials and can pose societal problems, for example when they grow in plumbing systems or on medical implants. Understanding the processes that promote the growth and disassembly of biofilms could lead to better strategies to manage these structures. We had previously shown that *Helicobacter pylori* bacteria are repulsed by high concentrations of a self-produced molecule, autoinducer-2 (AI-2) and that *H. pylori* mutants deficient in AI-2 sensing form larger and more homogeneously spaced biofilms. Here we used computer simulations of biofilm formation to show that local *H. pylori* behavior of repulsion from high AI-2 could explain the overall architecture of *H. pylori* biofilms. Our findings demonstrate that it is possible to change global biofilm organization by manipulating local cell behaviors, which suggests that simple strategies targeting cells at local scales could be useful for controlling biofilms in industrial and medical settings.

## Introduction

Bacteria often exist in aggregated, adherent communities called biofilms in which the cells are encased in a self-produced matrix and adopt distinctive three-dimensional architectures with heterogeneous cell spacing that gives rise to networks of channels. These biofilm structures confer increased resistance to environmental stressors such as antibiotics, changes in pH, and host immune defenses (1, 2). The architecture of mature biofilms contributes to their durability and resilience to perturbations by allowing for the flow of nutrients and waste products into and out of the cell aggregates (2–5). Biofilms create many commercial and biomedical problems for society, from biofouling of municipal waterworks to life-threatening infections by pathogens harbored on medical implants or the lungs of cystic fibrosis patients (1, 6, 7). Being able to understand and ultimately manipulate biofilm assembly and disassembly would help address several major industrial and biomedical challenges.

Biofilm assembly has been described alternatively as a developmental program controlled by stage-specific gene expression, similar to the development of a multicellular organism, or as the outcome of local adaptations of individual cells (8, 9). Distinguishing these two alternative possibilities is challenging because it can be difficult to discern whether biofilm phenotypes are achieved by optimizing group or individual fitness. For example, genes identified through forward genetic screens as being required for normal biofilm structures could be interpreted alternatively as being part of a biofilm genetic program or as controlling certain cellular behaviors that contribute to the self-assembly of biofilm structures. As a complement to experimental studies, computational modeling has played an important role in the study of biofilm assembly because it provides researchers with the opportunity to test and refine their understanding of the minimal set of parameters that can give rise to biofilm structures observed in nature (10–12).

A distinguishing feature of the biofilm lifestyle is that cells in close proximity can produce and respond to secreted molecular signals on small spatial and temporal scales. One such example of a secreted signal is the class of quorum-sensing molecules that serve as density-dependent forms of communication to influence group behaviors. These can include species-specific molecules, such as acylated homoserine lactones produced by many Gram-negative bacteria and perceived by species-specific receptors. Another example of a quorum-sensing molecule is the tetrahydroxy furan molecule autoinducer 2 (AI-2), which is produced by many bacteria through a common metabolic pathway but elicits different responses through species-specific receptors. Quorum-sensing molecules often regulate gene expression, including genes involved in biofilm growth and dissolution, by acting through canonical signal transduction pathways (13, 14). In this context, quorum-sensing molecules can be viewed as master regulators of biofilm developmental programs. AI-2 specifically has been shown to influence the overall structure of bacterial biofilms in diverse organisms such as *Bacillus, Streptococcus, Aggregatibacter, Pseudomonas, Escherichia, Vibrio, and Helicobacter* (15–22). In addition to regulating gene expression, AI-2 can elicit more immediate behaviors in bacteria through chemotaxis signal transduction that directs bacterial movement relative to a chemical gradient (16, 23–27). In the case of *Helicobacter pylori*, we showed that AI-2 is perceived as a chemorepellent (28), whereas *Escherichia coli* perceives AI-2 as a chemoattractant (25).

Previous experimental work from our group showed that both *H. pylori* biofilm mass and structural patterning are influenced by AI-2 chemotaxis. To determine the role of AI-2 in *H. pylori* biofilm formation, we constructed strains that were defective for AI-2 production *(luxS*^*-*^), AI-2 chemoreception (*cheA*^*-*^, *tlpB*^*-*^, *aibA*^*-*^, *aibB*^*-*^), or overproduced AI-2 (*luxS*^*OP*^). We measured the biomass of the resulting biofilms using a crystal violet assay. We also measured the structural heterogeneity of the resulting biofilms by imaging them with fluorescence microscopy and quantifying a lacunarity metric that captures morphological features such as roughness of biofilm edges and patchiness of surface coverage. We observed that both AI-2 sensing and production mutants formed larger biofilms with more homogenous organization, whereas the strain that overproduced AI-2 formed smaller, more heterogeneously structured biofilms (16).

Our experimental observations are consistent with a role for AI-2 chemorepulsion in shaping biofilm structure. For example, bacterial cells that chemotax away from AI-2 would be motivated to leave and deterred from joining a biofilm that is a concentrated source of AI-2. Our experimental results, however, could not rule out the possibility that additional functions of AI-2 signaling, such as regulation of global gene expression programs, contribute to the overall architecture of *H. pylori* biofilms. Here we used agent-based modeling to ask whether individual cellular behaviors of AI-2 production and chemotaxis are sufficient to produce global features of biofilm structures observed experimentally.

To explore the extent to which AI-2 chemotactic responses could explain our experimental observations, we used a well-established biofilm modeling platform, Individual-based Dynamics of Microbial Communities Simulator (iDynoMiCS) (29), which simulates behaviors of individual bacterial cells to understand larger, community behaviors. We implemented several critical modifications to iDynoMiCS in order to explore whether AI-2 chemotactic responses could recapitulate our experimentally observed biofilms. First, we expanded the models to include three spatial dimensions. Next, we included a population of planktonic (free swimming) cells that were continually introduced into the bulk medium and could join the biofilm. Additionally, cells from the biofilm could leave and become part of the planktonic population. Finally, we introduced AI-2 as a compound that was produced by individual cells as a function of their metabolic capacity and that diffused through the three dimensional space.

With the addition of AI-2 production and chemoreception to our modeling platform, we recapitulated our previous experimental data showing that biofilms of strains lacking the ability to produce or sense AI-2 were larger than wild type biofilms. In addition, the architecture of the biofilms, including spacing of cell groups within the biofilms, matched well between the experimental and modeled biofilms. Finally, our modeling of biofilms contributed new insight into the demographics dictating biofilm size, suggesting that cell dispersal is a major contributor to the reduced biofilm mass of AI-2-responsive versus non-responsive cells. These results indicate the utility of our modified iDynoMiCS platform for studying chemotaxis in biofilm dynamics and provide support for the view that local cellular behaviors of AI-2 chemotaxis can explain global features of biofilm formation and patterning.

## Materials and Methods

### Computational modeling of biofilms

The simulation of the growth of biofilms was accomplished using the agent-based modeling package iDynoMiCs. Individual cells are represented as discrete spherical agents with programmable behaviors that are subject to influence from other agents and their surrounding environments. The model consists of an evenly spaced grid of three dimensions with two compartments – the bulk and the bacterial. The bulk compartment at the top represents well-mixed bulk solutes that interface with the bacterial compartment at the bottom through a diffusion liquid boundary layer. Solutes are represented by concentration fields changing due to diffusion and from uptake by the cells in the bacterial compartment that provides a surface for initial seed cells to attach. As the cells uptake solutes, they can grow and divide or die above or below certain set size thresholds. These processes of growth and division lead to mechanical stress between the cells, which is relieved through a shoving algorithm. This shoving and the simulation of other physical forces on the biofilm dictate the formation of the biofilm’s structure.

To represent a bacterial population with both biofilm-attached cells and planktonic cells and to simulate the dynamics of cells joining and leaving a biofilm, we extended iDynoMiCs to include new agents with attributes and behaviors specific to planktonic cells. This version is available at https://github.com/alexwweston/iDynoMiCS. Cells can be either biofilm-associated or planktonic cells capable of movement in two or three dimensions. A set number of planktonic cells are introduced into the simulation from the bulk compartment at a chosen interval and removed from the simulation if they leave the boundaries of the bacterial compartment. An individual planktonic cell will move at a random distance between 0 and its maximum distance at a random angle. If it ends its movement within a certain distance from a biofilm-associated cell, it will then switch from planktonic to biofilm-associated behaviors. The maximum distance to move and the threshold distance for joining a biofilm are simulation parameters.

Planktonic cells are also given behaviors to simulate chemotaxis response. A chemotaxing planktonic cell has attributes from the solute it identifies as a chemoeffector, whether or not it exhibits an attractive or repellent response to this chemoeffector, and the threshold for recognizing this chemoeffector. Before moving, a planktonic cell will detect the concentration of its chemoeffector at its current location. If it is above its chemoeffector threshold, it will detect the gradient of the chemoeffector and move at an angle towards or away from this gradient depending on its response.

The attributes and behaviors of biofilm-associated cells are additionally extended to simulate biofilm-associated cells leaving the biofilm and becoming planktonic. A biofilm-associated cell has attributes for its chemoeffector, a threshold for recognizing this chemoeffector, and a probability for leaving the biofilm if this threshold is surpassed. At the end of each interval in the simulation, cells on the periphery of the biofilm will check the local concentrations of their chemoeffector. If the concentration is above its chemoeffector threshold, that cell has a chance of leaving the biofilm at a frequency equal to its leaving probability. Upon leaving the biofilm, that cell becomes a planktonic cell and moves from the biofilm at a random angle away from the chemoeffector gradient. The chosen chemotaxis threshold corresponds to the concentration of AI-2 at which planktonic cells contribute to the population of the biofilm at the midpoint between cells never joining the biofilm and cells always joining the biofilm (Supplemental Figure 1).

To model the production of AI-2 by the cells, we examined multiple AI-2 production regimes, each creating different concentrations and localization patterns of AI-2 in the model biofilms. These three regimes included: constitutive production of AI-2, production of AI-2 tied to the growth reaction, and production of AI-2 tied to the growth reaction with additional update of AI-2 by the cells. Although the distribution of the AI-2 molecule within a biofilm is unknown, we measured the total concentration of AI-2 at different time points in *H. pylori* biofilms and compared these results with the total concentrations of AI-2 generated in our simulations under the different regimes (data not shown). From these results, we chose to model AI-2 production where it was tied to the growth reaction with uptake by the cells.

### Setting up and running simulations

We simulated the movement of bacteria through a 280x280x280 µm space for a period of 24 h. The space was modeled as a 33x33x33 grid. Fluid movement was simulated using a major time step size of 1.0 h, and bacterial behaviors (movement, joining, leaving) were updated at minor time steps of 0.05 h. Each simulation was seeded with 100 bacteria cells randomly placed on the bottom layer of the simulated grid. Outputs for visualization were recorded at the end of every major time step. Other parameters for concentration and diffusivity of solutes and the cell attributes of agents were taken from measurements of *E. coli* biofilms used in other simulations under iDynoMiCs. Erosion and sloughing processes that can be modeled in iDynoMiCs were turned off for these simulations. A full list of these parameters that were static in our simulations is available in Supplementary Table 1.

Parameters that were introduced in this new model were tested across a wide variety of ranges and values were chosen where moderate behavior was observed. Microbial growth kinetics were modeled using the Monod growth equation with an additional term representing the production of AI-2. The values for these parameters and equations for the wild type strain used in the simulations are found on Supplementary Table 2. Mutant strains used in the simulations use minor modifications of these values, which are highlighted on Supplementary Table 3. The mutant chemotaxis strain is given an infinite value for its chemotaxis threshold causing it to never detect its chemoeffector, the mutant overproducer strain is given a larger AI-2 yield coefficient, and the mutant strain defective in AI-2 production creates an arbitrary alternative product other than AI-2 from its growth reaction.

### Visualizing the biofilms

The visualization of the agent-based simulation of gut microbes was created using custom-built codes developed in C++, using OpenGL for the graphics and Qt for the user interface. The simulations are run within iDynoMiCS, which exports the entirety of the simulation in XML. Each microbe is displayed as a sphere that has a radius dictated by the simulation and a color based on the microbe type, and in some scenarios, modified based on family, genealogy, generation, or birthday. Each founding cell is labeled in a different shade of pink and the daughter cells remain the same color as the original founding cell to allow for recognition of clones. The code is open source and can be downloaded at https://bitbucket.org/kpotter/vizr.

The visualization of the AI-2 gradients via contours was done using the R library filled.contours. To create these images, the data is loaded into R, a single slice of the data volume is extracted at a specified timepoint, and this data is used as input to the contours function. The R code is provided in Supplemental Materials and Methods.

### Calculating lacunarity

The simulated biofilms all become 100% confluent by 24 h. To compare more directly to the experimental biofilms, which were often not confluent by 24 h, a bottom portion of each simulated biofilm was removed. To decide how much to trim off, the percent cell coverage across all experimental wild type images was determined to be approximately 43% percent coverage using ImageJ. Removing the bottom 98 µm from the simulated biofilms resulted in 43% coverage for a representative set of wild type biofilms, viewed from the top down (Figure 4A). Therefore, 98 µm was removed from all simulated biofilms and lacunarity was determined. To determine the lacunarity score, we opened the experimental or trimmed simulated biofilm images in ImageJ, converted them to black and white, adjusted the threshold to a set cutoff, and analyzed the resultant images using the FracLac plugin.

**Figure 4.**
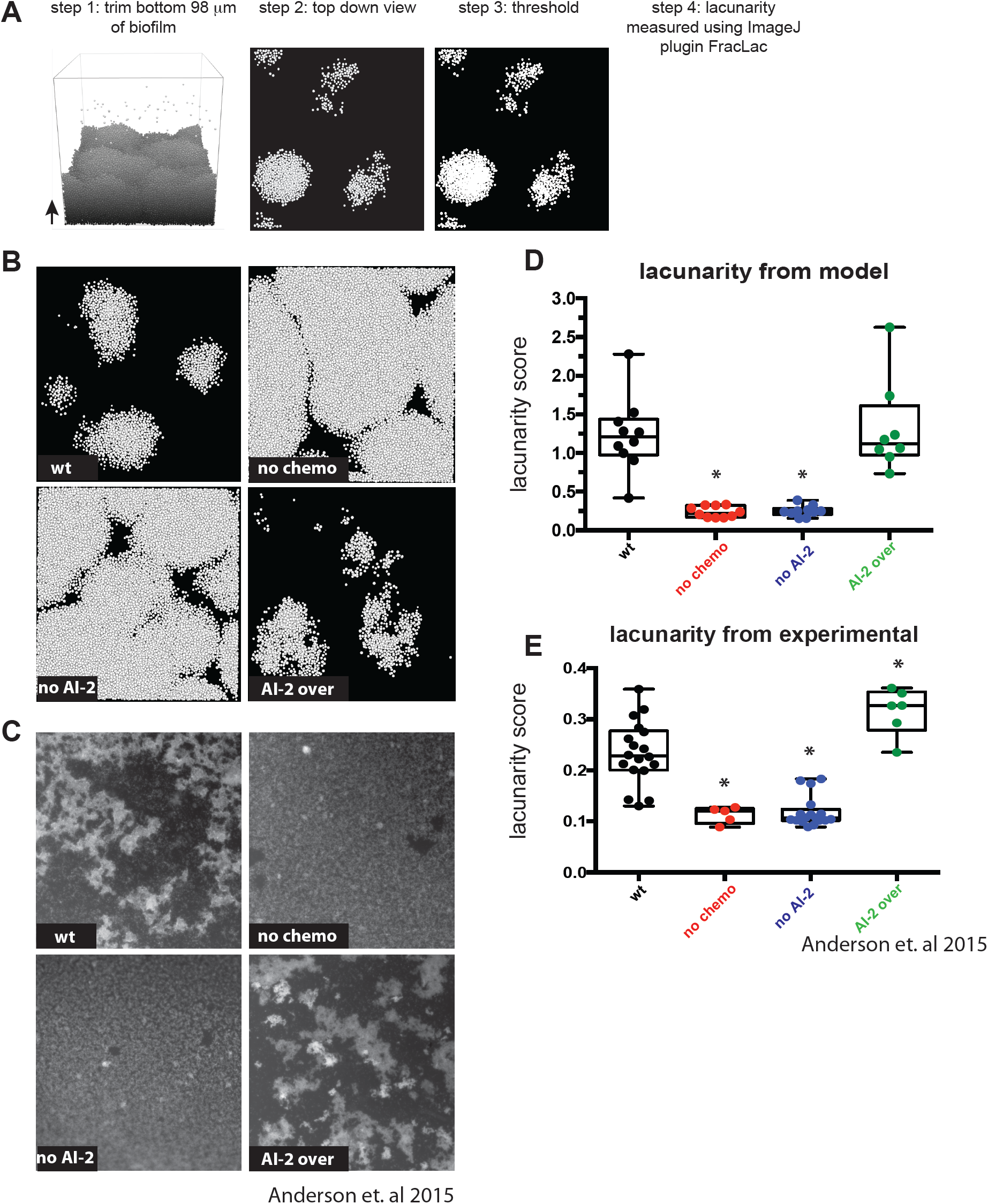
Modeling confirms AI-2 chemotaxis and production influence biofilm organization. A) Lacunarity analysis pipeline for the modeled biofilm images. Bottom 98 µm removed from each 24 h biofilm across all genotypes (see Materials and Methods). The top-down view is used to be able to compare to the experimental images (4C). Using ImageJ, each image was thresholded and then run through FracLac to determine the lacunarity score. More details can be found in the Materials and Methods section. B) Example images of all four modeled genotypes from the top-down. C) Example images of experimental *H. pylori* biofilms grown on glass slides, from Anderson et al. 2015. D) Lacunarity scores graphed for modeled biofilms (n = 8-10). E) Lacunarity scores graphed for each experimental biofilm for each genotype from Anderson et al. Stars in D and E indicate a significant difference from wild type, results determined using a one-way analysis of variance (p < 0.05). 4E data from Anderson et al. 2015 (16).

## Results

### Addition of chemotaxis and AI-2 production to agent-based modeling of biofilm formation

Agent-based models are useful tools for exploring how simple interactions between cells contribute to the overall properties of bacterial communities, such as biofilms. iDynoMiCS simulates biofilm formation by taking into account biologically relevant parameters such as nutrient concentrations, nutrient diffusion rates, and cell division and spacing [see Supplementary table 1 for parameters used in the simulation (29)]. To investigate the role of AI-2 mediated chemotaxis in biofilm architecture, we extended the iDynoMiCS model by introducing several properties, including: three-dimensional chemotaxis, planktonic (free swimming) cells, cells joining and leaving the biofilm, and AI-2 production. These new iDynoMiCS additions were critical for exploring how chemoreception of AI-2 shapes *H. pylori* biofilms. In addition, these developments necessitated a new visualization tool that aided in data interpretation (see Materials and Methods).

Our extended model starts with 100 bacterial cells randomly placed on the two-dimensional surface at the bottom of a container that is continually supplied with fresh, nutrient-containing medium. These cells expand and proliferate according to the iDynoMICS growth and spacing algorithms. We allowed new planktonic cells to enter the container throughout each 24 h simulation (Figure 1A and Supplementary movies). The planktonic cells moved through the space according to a chemotaxis algorithm (see Materials and Methods). Planktonic cells would join the simulated biofilm if they swam close enough to the biofilm surface and if the concentration of a chemorepellant was below a set threshold. In addition, cells at the biofilm edge could leave and enter into the planktonic pool. In our simulations, we chose the AI-2 chemotaxis threshold to be that at which planktonic cells contributed to the population of the biofilm to the extent that was defined as halfway between cells never joining the biofilm and cells always joining the biofilm (Supplemental Figure 1).

**Figure 1.**
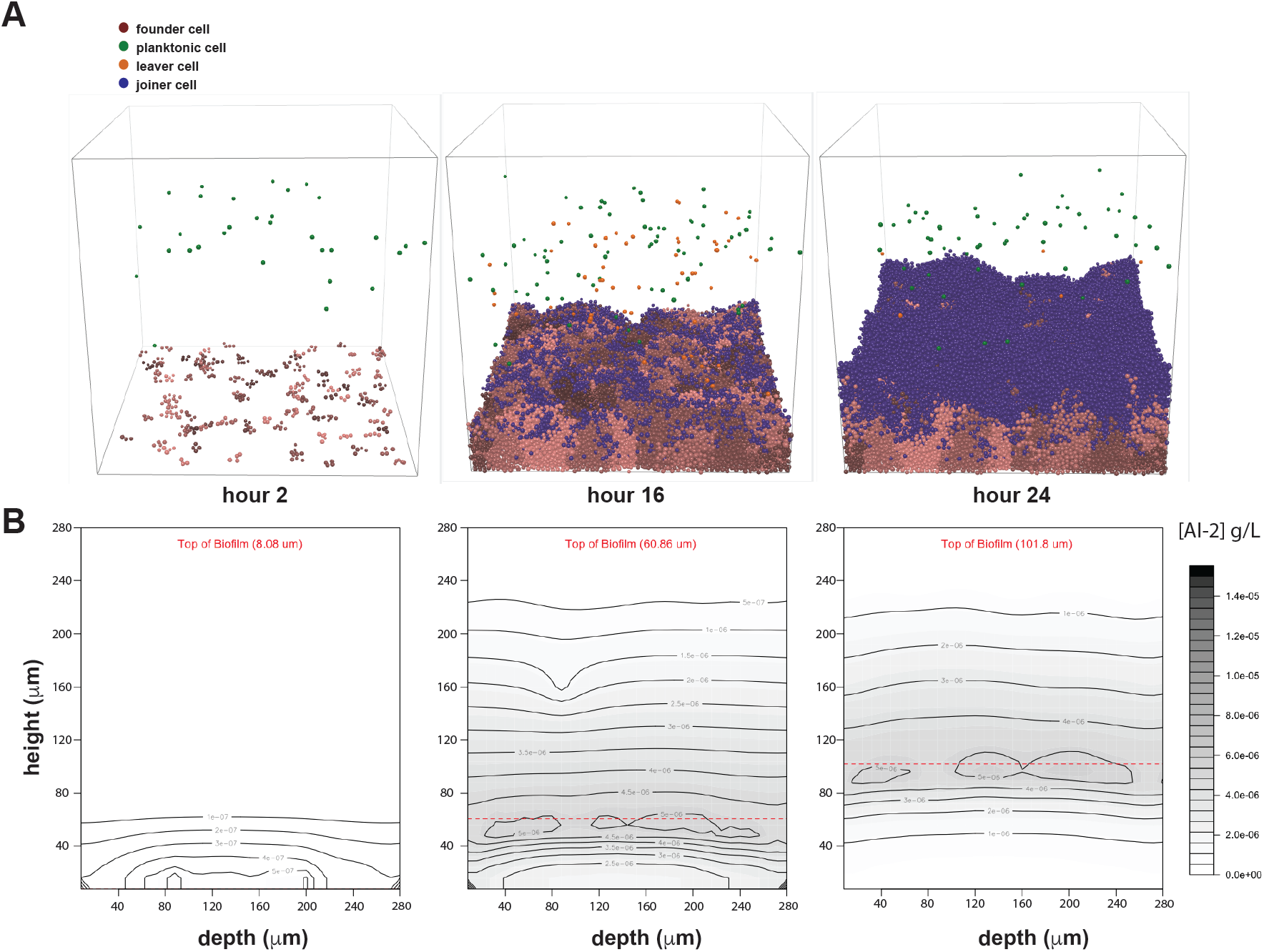
Time steps and AI-2 gradients of example wild type iDynoMiCS modeled *H. pylori* biofilm. A) Wild type biofilm after 2, 16 and 24 h of growth. Each sphere represents a modeled bacterial cell with colors corresponding to different cell behaviors (see legend). Note there is a mix of cells leaving, dividing from the original founding population and cells joining the biofilm. Each grouping of pink cells represents a clonal population. B) Shown are corresponding AI-2 concentration graphics below each time point shown in A. The AI-2 concentration is a representative vertical slice through the center of the 3D modeled biofilm, with darker color representing higher concentrations of AI-2.

After testing several models of AI-2 production (see Materials and Methods), we chose a model that tied AI-2 production directly to the growth and metabolism of each cell or agent. This model is reasonable because AI-2 is produced as a by-product of the activated methyl cycle (30). We also incorporated into the model a constant cellular uptake of AI-2, which is common in bacteria (31). We do not yet know whether *H. pylori* has an active AI-2 uptake mechanism, but incorporating a constant uptake parameter best recapitulated our experimental measurements of AI-2 [(16, 23) and data not shown]. In the iDynomics simulations, cells near the surface of the modeled biofilms have more access to fresh nutrients and therefore divide and produce AI-2 at a higher rate than cells in the middle or bottom of the biofilm (Figure 1B). The constant cellular uptake of AI-2 in the model resulted in a lower concentration of AI-2 in the volume just below the surface of the biofilm (Figure 1B).

### Modeling recapitulates biofilm mass as a function of AI-2 chemorepulsion

Using the model, we tested whether we could recapitulate the outcomes of our previous experiments demonstrating an important role for AI-2 production and chemorepulsion in *H. pylori* biofilm mass and patterning (16). To simulate these experiments, we modeled the strains and conditions used in this experimental work. The strains included wild type cells, cells unable to chemotax, cells unable to produce AI-2, and cells that overproduce AI-2. For each of these genotypes we ran 30 individual iterations and compared the number of cells in our simulated *H. pylori* biofilms to those of experimental work (Figure 2). Wild type cells produced moderately sized biofilms in both the model and the experimental set-up, while cells that could not produce AI-2 or chemotax away from AI-2 produced larger biofilms. Finally, both experimental and modeling results revealed that the AI-2 overproducer made smaller biofilms. This data served as confirmation that our modeling platform could recapitulate experimental results.

**Figure 2.**
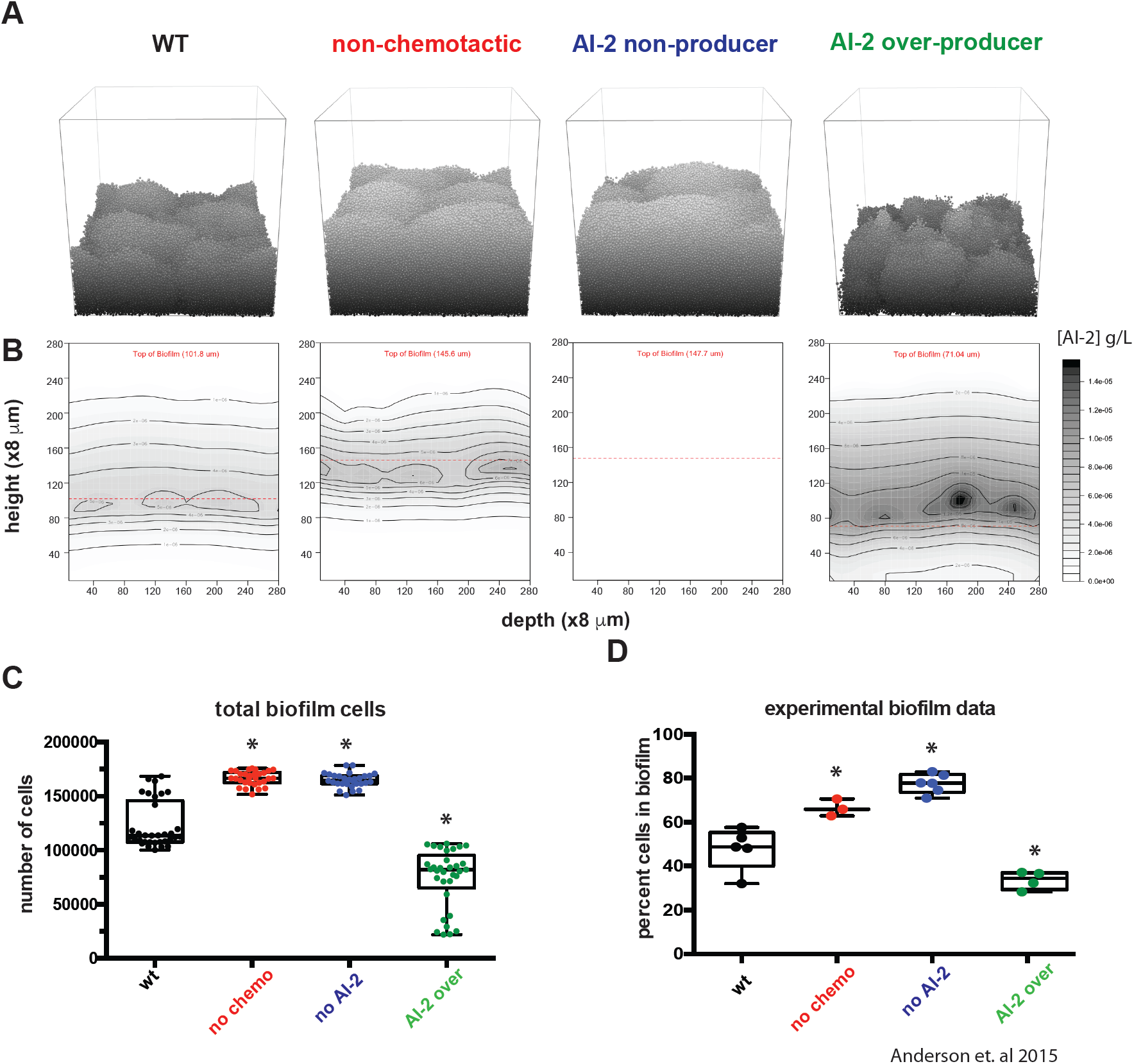
Modeling confirms AI-2 chemotaxis and production alter overall biofilm size. A) Representative images of 24 h biofilms for each of the four strains in grayscale to show contours. To simplify, only the founding population, their progeny and joiners are shown. Planktonic cells have been removed for simplicity. B) The associated AI-2 gradients for panels in A. C) Total number of cells attached to the modeled biofilms at timepoint 24 h. n = 30 D) The size of the experimental biofilms from Anderson et al. are graphed according to percentage of cells in the biofilm (compared to planktonic). Stars indicate a significant difference from wild type. Statistics for C and D were determined using a one-way analysis of variance (p < 0.05). 2D data from Anderson et al. 2015 (16).

### Modeling predicts subcellular populations that contribute to biofilm mass

The model afforded us the opportunity to examine the cellular demographics contributing to biofilm mass, which would be difficult to do experimentally. We modeled biofilm formation for 24 h in 30 parallel simulations and tallied the individual leaving events and joining events. Wild type biofilms showed equivalent numbers of joining and leaving cells (Figure 3). As expected, biofilms of non-chemotactic and AI-2 non-producing cells had no leaving events, since AI-2 chemorepulsion was the only leaving mechanism in the model. The AI-2 over-producer strain had a dramatic increase of leaving events, which was expected given the higher concentration of AI-2 near the surface of the biofilm (Figure 2B) that would drive cells to chemotax away from the biofilm.

**Figure 3.**
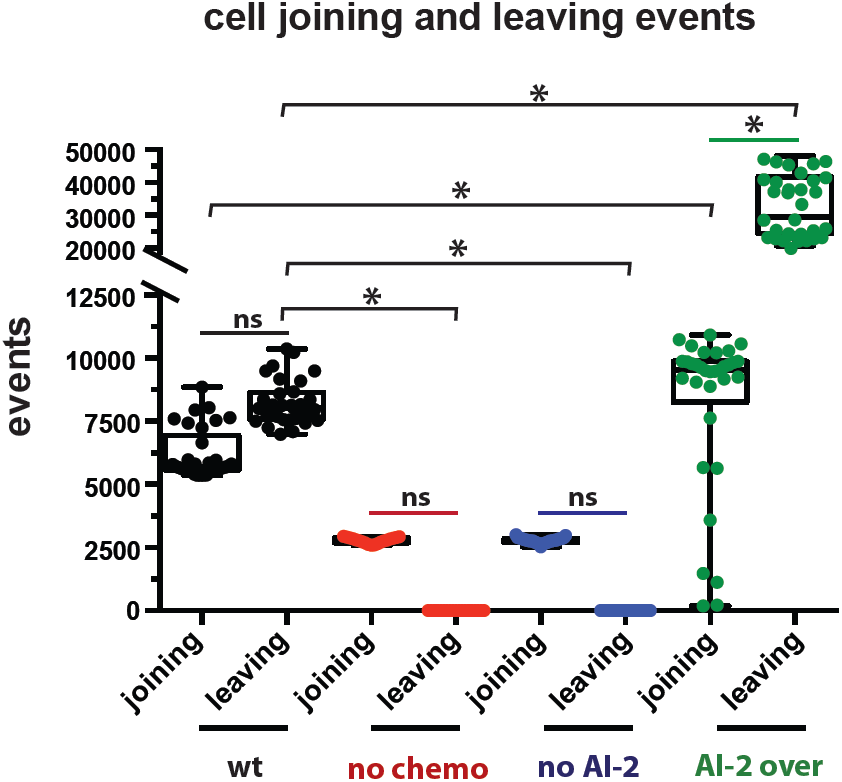
Modeling confirms AI-2 chemotaxis and production influence the behavior of the cells in the biofilm. Each leaving and joining event from 0-24 h of the modeled biofilms was graphed by genotype. Stars indicate a significant difference, results determined using a one-way analysis of variance (p < 0.05). n= 30 biofilms

Interestingly, both the non-chemotactic and AI-2 non-producers had a reduction in the number of joining events as compared to the wild type population, despite not experiencing chemorepulsion from the biofilm. Also counter-intuitively there were more overall joining events in the AI-2 over-producer strain biofilm than in the wild type population. As discussed below, the number of joiners could be explained by the differences in architectures and specifically surface areas and joining opportunities afforded by the growing biofilms of the different strains. Overall, our modeling supports the idea that AI-2 chemorepulsion promotes a balance of leaving and joining events that influences the global biofilm size.

### Modeling recapitulates the impact of AI-2 chemorepulsion on biofilm spatial organization

Finally, we used modeling to confirm that AI-2 shapes *H. pylori* biofilm architecture. We had shown previously that we could quantify the heterogeneity of biofilms using a lacunarity metric, which measures spacing between patterns and boundary smoothness. Experimental biofilms were grown on glass slides, fixed and stained with DAPI and visualized with epifluorescence. The cellular component of the biofilm was defined using an intensity threshold and the resulting images were analyzed using an ImageJ plugin, FracLac, to quantify lacunarity [Figure 4, (32–34)]. We took 10 simulated biofilms for each genotype and performed a similar analysis of a top view of the structure (Figure 4A).

Visually, the simulated biofilm structures (Figure 4B) resembled the experimental data (Figure 4C). The wild type and AI-2 overexpressing strains produced biofilms with marked spacing between cell patches, whereas the non-chemotactic and AI-2 non-producing strains formed much more homogeneous structures. Plotting the resulting lacunarity scores revealed a striking similarity between experimental and modeling data (Figure 4D and E).

## Discussion

In this study, we used agent-based modeling to explore whether local cell chemotaxic responses to a self-produced molecule could explain biofilm growth and patterning properties. By extending the iDynoMiCS modeling platform to include three-dimensional chemotaxis, cell joining and leaving events, and AI-2 production, we were able to recapitulate our experimental observations of *H. pylori* biofilm formation with a collection of strains with different AI-2 production and perception properties (16). We showed that cells unable to make or chemotax away from AI-2 produced larger biofilms than wild type cells. These biofilms also differed in their organization with more homogenous cell spacing and smaller gaps between cell clusters. Over-production of AI-2 resulted in smaller and more heterogeneously spaced biofilms. Mature biofilms are complex structures with towers and channels that facilitate fluid flow for efficient oxygen and nutrient permeation, waste excretion, and cell turnover (2–5). We found that by modeling local chemotactic responses to a self-produced molecule, we could simulate the assembly of biofilms with the global property of high lacunarity, characteristic of biofilms with extensive channels (Figure 4). The agreement between our simulations and experimental results supports the idea that local cellular behaviors, such as production and chemoavoidance of AI-2, can explain global architectural features of bacterial biofilms.

Our modeling approach allowed us to dissect the demographics of biofilm assembly in a way that would be difficult to do experimentally without sophisticated genetic tools for marking cell lineages. As expected, in our model cells left the biofilm when they were programmed to chemotax away from AI-2, and they left in greater numbers when the biofilm cells produced more AI-2. We did not initially expect the wild type and AI-2 overproducer populations to have more cells join the biofilms than the populations without chemotaxis or AI-2 production. However, inspection of the biofilm structures assembled in these different models showed that the wild type and overproducer biofilms had many more gaps and edges, creating more extensive surface area that planktonic cells would stochastically encounter and then join at a certain probability. This difference in surface area and joining opportunities could explain the higher numbers of joiners in the populations of cells engaging in AI-2 chemorepulsion. In addition, the heterogeneous architectures of these biofilms would create local minima in AI-2 concentrations and opportunities for joining even in the context of AI-2 chemorepulsion. Differences in the local AI-2 concentration landscapes could explain the higher number of joiners seen with the AI-2 overproducer versus wild type cell populations.

Although our model recapitulated several features of AI-2 dependent biofilm assembly observed experimentally, it is based on certain assumptions about AI-2 fluxes that are likely to be oversimplifications. In our current model, AI-2 production is linked to metabolic activity and uptake is constant. When examined experimentally, parameters of AI-2 production, uptake and sensing are known to vary greatly between bacterial species and depending on cells’ metabolic states (35–38). Using deterministic simulations of AI-2 production from a system of ordinary differential equations, Quan and colleagues showed that variability in AI-2 uptake within a modeled biofilm can lead to desynchronization of autoinduction across the community (39), highlighting the importance of considering heterogeneities in AI-2 fluxes. In addition, AI-2 could be produced from sources other than the bacterial constituents of a biofilm. For example, mammalian host tissues were recently shown to synthesize an AI-2 mimic that is sensed by bacterial AI-2 receptors (40). Future iterations of the model could incorporate more detailed parameters of AI-2 fluxes, but these would need to be tailored to the specific bacterial species and environments being modeled.

Most bacteria exist not in mono-cultures but rather in multi-species consortia (41). AI-2 is known to contribute to the organization of such consortia, for example in biofilms that assemble on the enamel surfaces of teeth (42, 43). Recently, Laganenka and Sourjik showed that in a simple two-member biofilm community of *Enterococcus faecalis* and *Escherichia coli* cells, AI-2 chemotaxis plays an important role in biofilm growth and patterning. In this model community, both species produce AI-2 but only *E. coli* chemotaxes toward it (27). It would be interesting to apply our modeling approach to this experimental system to test whether it would recapitulate observed architectural features, such as the spatial segregation of *E. faecalis* and *E. coli* cells. More generally, by applying our modeling approach to complex multi-species communities and assigning simple AI-2 production, chemoattraction, and chemorepulsion behaviors to different members, one could explore the extent to which local AI-2 chemotactic responses could explain global spatial patterning observed in multi-species communities.

## Acknowledgements

Research reported in this publication was supported by the National Institute of Diabetes and Digestive and Kidney Diseases and the National Institute of General Medical Sciences of the National Institutes of Health under award numbers R01DK101314 and P50GM098911 (to K.G.) and the Medical Research Foundation Oregon Scientist Development Award (to E.G.S). The content is solely the responsibility of the authors and does not necessarily represent the official views of the National Institutes of Health.

